# Simple ways to avoid collecting highly ^137^Cs-contaminated *Aralia elata* buds

**DOI:** 10.1101/2023.09.18.558164

**Authors:** Masaru Sakai, Mirai Watanabe, Masami Kanao Koshikawa, Asuka Tanaka, Akiko Takahashi, Seiichi Takechi, Mai Takagi, Takashi Tsuji, Hideki Tsuji, Toshimasa Takeda, Jaeick Jo, Masanori Tamaoki, Seiji Hayashi

## Abstract

Collection and cooking of wild vegetables have provided seasonal enjoyments for Japanese local people as provisioning and cultural ecosystem services. However, the Fukushima Daiichi Nuclear Power Plant accident in March 2011 caused extensive radiocesium contamination of wild vegetables. Restrictions on commercial shipments of wild vegetables have been in place for the last 10 years. Some species, including buds of *Aralia elata*, are currently showing radiocesium concentrations both above and below the Japanese reference level for food (100 Bq/kg), suggesting that there are factors decreasing and increasing the ^137^Cs concentration. Here, we evaluated easy-to-measure environmental variables (dose rate at the soil surface, organic soil layer thickness, slope steepness, and presence/absence of decontamination practices) and the ^137^Cs concentrations of 40 *A. elata* buds at 38 locations in Fukushima Prefecture to provide helpful information on avoiding collecting highly contaminated buds. The ^137^Cs concentrations in *A. elata* buds increased significantly with increases in the dose rate at the soil surface. Meanwhile, the ^137^Cs concentration in *A. elata* buds were not reduced by decontamination practices. These findings suggest that measuring the latest dose rate at the soil surface at the base of *A. elata* plants is a helpful way to avoid collecting buds with higher ^137^Cs concentrations and aid in the management of species in polluted regions.

## Introduction

Collecting and cooking wild vegetables (*sansai*) provides seasonal enjoyment for Japanese people and such traditional activity promotes communication among local people through sharing of dishes and information [1]. However, the Fukushima Daiichi Nuclear Power Plant accident in March 2011 resulted in radionuclide contamination, leading to the termination of such activities [2, 3]. Collecting various wild vegetables is restricted in some contaminated areas due to radiocesium concentrations exceeding the Japanese reference level (100 Bq/kg) [4, 5]. Among the vegetables, the buds of Koshiabura (*Eleutherococcus* [*Chengiopanax*] *sciadophylloides* [Araliaceae]), which is known as the “Queen of *sansei*,” exhibits particularly high radiocesium concentrations compared to other wild vegetables [6–8]. Consequently, commercial shipping of this species is restricted in 113 municipalities in Japan [9].

Radiocesium concentrations in several wild vegetable species have decreased over the past decade [10] and some species are exhibiting concentrations below the reference level. For example, the reported ^137^Cs concentrations in wild vegetables collected in Iitate village, one of the former evacuation zones, include a substantial number of minimally contaminated species [11]. Among them, the buds of Taranoki (*Aralia elata* [Araliaceae]), which is one of the most popular wild vegetables in Japan known as the “King of *sansai*,” exhibited ^137^Cs concentrations that varied widely above and below the reference level (Fig 1).

**Fig 1.**
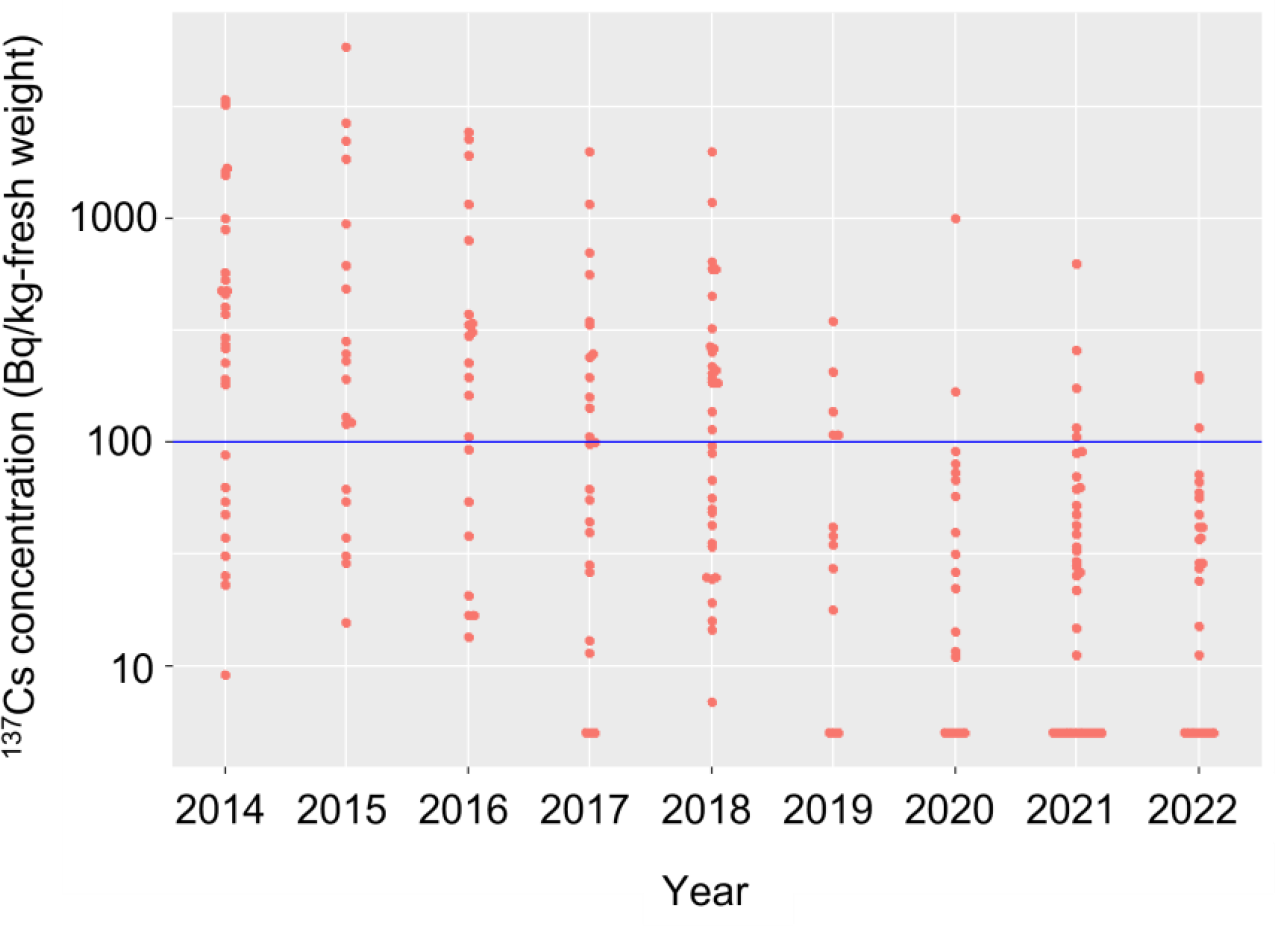
^137^Cs activity concentrations in *Aralia elata* buds collected in Iitate village, Fukushima from 2014 to 2022. The blue horizontal line indicates the Japanese reference level for foods (100 Bq/kg). Data were retrieved from Iitate village (2022) [11]. ^137^Cs activity concentrations below the detection limits were regarded as 5 Bq/kg. Reported ratios of ^134^Cs to ^137^Cs outside the range of 0.8–1.2 on 11 March 2011 were excluded to ensure reliability.

*Aralia elata* is a deciduous shrub species distributed in East Asia, including the Japanese Archipelago, the Korean Peninsula, China, and Russia. As a typical pioneer species, *A. elata* primarily inhabits sunlit spaces on forest edges, roadside slopes, and riparian zones. The buds of this species (Taranome) are typically collected during spring and are often cooked as tempura and ohitashi in Japan or boiled as medicine in Korea and China. Forty-three Japanese municipalities restrict commercial shipments of wild *A. elata* buds; only Koshiabura buds are subjected to more stringent limitations [9]. However, as mentioned above, ^137^Cs concentrations in *A. elata* buds vary widely, including values below the reference level, even within a single municipality (Fig 1). This trend suggests that potential factors are decreasing and/or increasing the ^137^Cs concentrations in *A. elata* buds. For example, the ^137^Cs inventory in the soil (or the dose rate at the soil surface) is a fundamental factor determining the ^137^Cs concentrations in various organisms, including plants [12–14]. Moreover, *A. elata* predominates in open spaces that often lie near areas of human activity, such as roadside slopes and clear-cut areas. Soils in these areas may have undergone decontamination efforts following the Fukushima accident [15]. Forested areas have rarely been decontaminated; thus, leaf abscission and circulation of ^137^Cs in the organic soil layer may continue to deposit bioavailable ^137^Cs [16]. Furthermore, the steepness of the slope can affect the ^137^Cs concentration in *A. elata* buds due to the pronounced effects of soil erosion on steeper slopes, which reduces the amount of deposited ^137^Cs in the soil [17].

As collecting and cooking *A. elata* buds have provided seasonal enjoyment for local people, proposing easy ways to avoid obtaining highly contaminated buds would be helpful. In this study, we collected *A. elata* buds and constructed an environmental variable dataset around the *A. elata* plants that can be easily obtained using commercially available equipment or public data. This investigation was performed in six municipalities of Fukushima Prefecture with a variety of initial ^137^Cs deposition levels. The environmental variable dataset included the dose rate at the soil surface, organic soil layer thickness, slope steepness, and presence/absence of a decontamination practice. We assessed which environmental variable(s) should be avoided to collect highly contaminated buds. Simply identifying these factors could help revive traditional wild vegetable collection and cooking practices, as well as aid decontamination programs. This is particularly crucial to preserve the traditional *sansai* culture within contaminated regions.

## Methods

### Study sites and bud collection

This study was conducted at 38 sampling locations in Iitate and Katsurao villages, Kawamata town, and Minamisoma, Nihonmatsu, and Tamura cities in Fukushima Prefecture, Japan (Fig 2). The initial ^137^Cs deposition after the Fukushima Daiichi Nuclear Power Plant accident varied widely, ranging from 41 to 1,200 kBq/m^2^ in the study locations [18]. The habitats of *A. elata* plants sampled in this study were categorized as forest, forest edge, or roadside slope, which are preferred habitats for this species. The height of the 38 sampled plants was approximately 2.0 m. Because *A. elata* generally grows 20–60 cm/year, we assumed that the sampled plants had not been contaminated by the initial deposition on their surfaces in 2011. While *A. elata* generates underground stems for vegetative propagation, the plants we selected were treated as separated genets due to the significant distance between their habitats. Buds were collected from all plants from April to May in 2022 and 2023 to measure ^137^Cs concentrations. A total of 40 buds were collected, including four buds from two plants collected in 2022 and 2023 (one bud per plant for each year) and 36 buds from each of the remaining 36 plants in 2022 or 2023.

**Fig 2.**
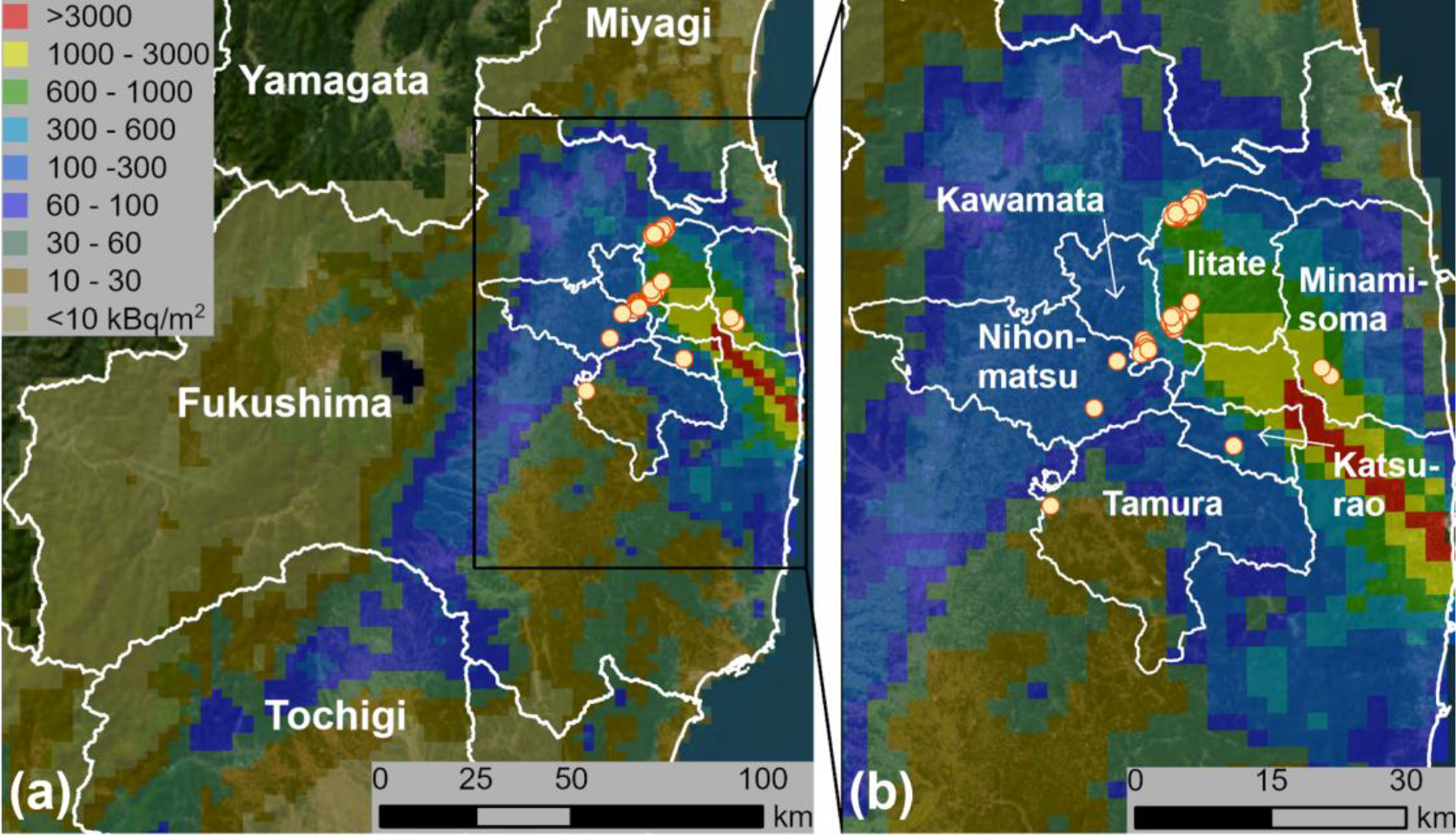
(a) Map of Fukushima and neighboring prefectures and (b) a close-up view of the study sites. The study sites are shown as circles. ^137^Cs deposition are shown using the data from the Fifth Airborne Radiation Monitoring by the Ministry of Education, Culture, Sports, Science, and Technology (MEXT) in 2012 [18]. The map is based on data obtained from Esri, Maxar, Earthstar Geographics, and the GIS User Community.

### Environmental variables

To identify “easy-to-measure” factors that increased the ^137^Cs concentrations in *A. elata* buds, we investigated the presence/absence of decontamination practices, the dose rate at the soil surface, slope steepness, and the thickness of the organic soil layer at the base of each *A. elata* plant. According to information gathered from the decontamination records provided by the Ministry of the Environment, the sampling locations included 24 decontaminated and 14 abandoned habitats.

Because several years have passed since decontamination, the current soil ^137^Cs inventory, particularly within the decontaminated areas, could change owing to recontamination from the surrounding environment [19]. To address this concern, we measured the dose rate at the soil surface at the base of each *A. elata* plant to serve as a reliable indicator of the ^137^Cs inventory (see *Aggregated transfer factors* below for details). The dose rates at the soil surface were determined using a scintillation survey meter (TCS-172B; Hitachi Aloka Medical, Tokyo, Japan). The sensor of the survey meter was placed at the soil surface, and thus the measured dose rates were assumed to be primarily affected by radiation from the soil, rather than that from the surrounding environment. Dose rate can also be measured using a portable dosimeter, which is generally available to the public, and thus we included this as an easy-to-measure variable.

Slope steepness was determined using the “Measure” iPhone application (iPhone 13; Apple Inc., Cupertino, CA, USA). We assumed that slope steepness could affect the soil runoff rate, which, in turn, is related to decreasing ^137^Cs content in the soil [17]. The cumulative thickness of the litter, fermentation, and humus layers at the soil surface was measured using a 3-point folding scale in the vicinity of each *A. elata* plant because the organic soil layer possessed abundant bioavailable ^137^Cs that has the potential to contaminate *A. elata* buds [16].

### Aggregated transfer factors

Estimating the potential internal radiation exposure after consuming contaminated wild vegetables is important for radiation protection [20]. This is particularly important to improve the quality of life of local people who wish to enjoy collecting and consuming wild vegetables [21]. In this regard, the aggregated transfer factor (T_ag_) is widely used to indicate the relationship between radionuclide concentration in wild vegetables and the soil. T_ag_ is expressed as the radionuclide activity concentration in wild vegetables (Bq/kg) divided by the radionuclide inventory in the soil (Bq/m^2^). Because accumulating T_ag_ values from spatiotemporally wide areas is helpful to understand the ^137^Cs dynamics in wild vegetables and predict the revival of *sansai* culture, this study also calculated the T_ag_ values based on the ^137^Cs activity concentrations in *A. elata* buds and ^137^Cs deposition in the soil that was estimated using a previously published method [22].

We first measured the dose rate at the soil surface and collected soil samples from 78 measurement points from 2020 to 2022 before investigating the *A. elata* buds. The 78 locations included forests, forest edges, and roadside slopes around Fukushima Prefecture (also three locations from the *A. elata* investigations). The dose rate at the soil surface was determined as described above. Then, the organic layer and the soil layer at depths from 0 to 10 cm were collected at each measurement point because these layers contain most of the ^137^Cs derived from the disaster [23]. The ^137^Cs soil inventory (kBq/m^2^) was estimated based on soil mass per unit area and the ^137^Cs activity concentrations of the collected samples. The procedures for measuring ^137^Cs activity concentrations are detailed below. A linear model was constructed based on the log_e_-transformed data for the soil surface dose rates and the ^137^Cs inventories in the soil using R 3.6.3 [24]. It was used to transform the measured dose rate at each *A. elata* plant into the ^137^Cs inventory at each point to calculate T_ag_ for the *A. elata* samples.

### Radioactivity measurements

The soil samples used to estimate the ^137^Cs inventory were dried at 25°C for at least 1 month, sieved through 2 mm mesh, and packed into 100 mL plastic containers. The dried soil weight was calculated based on the weight loss after subsequent drying of several grams of the samples at 105°C for at least 24 h. All fresh buds were gently washed with tap water, wiped, weighed, dried at 60°C for at least 2 days, and reweighed. The dried samples were finely ground using an electrical mill and packed into 100 mL plastic containers. The dry weights and the densities of the soil and bud samples were measured before the radioactivity measurements. The ^137^Cs activity concentrations in the soil and bud samples from all *A. elata* plants were determined by gamma-ray spectroscopy. Gamma-ray emissions were measured at an energy of 661.6 keV using a coaxial high-purity germanium detector system (model GC 2020; Canberra Japan, Tokyo, Japan). The accuracy of ^137^Cs activity was within 5% (error counts/net area counts). Sample activity was corrected for radioactive decay at the time of collection. After determining the ^137^Cs activity concentrations based on the bud dry weights, fresh weight concentrations were calculated based on water content estimated from the initial weight measurements (mean = 87.4%). The detector system was routinely calibrated using standard and blank samples, and tested annually by the manufacturer using reference samples to ensure measurement accuracy.

### Statistical analyses

A linear mixed model (LMM) was employed to identify the easy-to-measure factor(s) that affected the ^137^Cs activity concentrations in *A. elata* buds. Correlations between all pairs of environmental variables (dose rate at the soil surface, mean organic soil layer thickness, and slope steepness) were analyzed using Pearson’s correlation coefficient analysis; highly correlated variables (*ρ* > 0.7) were excluded from the models to avoid multicollinearity. The ^137^Cs activity concentrations in *A. elata* buds and the dose rates at the soil surfaces were log_e_-transformed to normalize the distributions and standardize the variance before constructing the model. A Gaussian error structure was used for the response variables. The sampling year and identifiers for each *A. elata* plant were included as random terms in the LMM to consider potential variabilities between years and among plants. The LMM was constructed using the lmer function of the lme4 package [25] in R 3.6.3 [24]. To visualize the relationship between ^137^Cs concentrations in buds and a single explanatory variable, the other explanatory variables were fixed at their mean values using the ggeffects R package [26]. The effect of the presence/absence of decontamination practices on ^137^Cs activity concentrations in *A. elata* buds was tested using analysis of covariance (ANCOVA), with presence/absence of decontamination practices as a factor and initial ^137^Cs deposition estimated from the Fifth Airborne Radiation Monitoring [18] as the covariate. The ^137^Cs activity concentrations in the buds were log_e_-transformed to normalize the distribution and standardize variance before ANCOVA. The analysis was performed using R 3.6.3 software.

## Results

### ^137^Cs concentrations in A. elata buds and the environmental variables

The ^137^Cs activity concentrations in *A. elata* buds ranged from 1 to 6,280 Bq/kg fresh weight. The values of the easy-to-measure environmental variables were dose rate at the soil surface, 0.10–6.50 μSv/h (Fig 3a and Table S1); mean organic soil layer thickness, 0–10.5 cm (Fig 3b and Table S1); and slope steepness, 3–49° (Fig 3c and Table S1). Because there was no pair of highly correlated explanatory variables with *ρ* > 0.7, all three environmental variables were used as explanatory variables in the LMM. The LMM indicated that the dose rate at the soil surface had a significant positive effect on the ^137^Cs activity concentrations in *A. elata* buds (Table 1; Fig 3). The ^137^Cs activity concentrations in *A. elata* buds showed a weak positive relationship with organic soil layer thickness and a negative relationship with slope steepness, but the relationships were not significant. The presence/absence of decontamination practices did not affect the ^137^Cs activity concentrations in *A. elata* buds (*t* = –1.124, *P* = 0.268), but the initial ^137^Cs deposit had a significant positive effect (*t* = 7.552, *P* < 0.001), as illustrated in Fig 4. These results indicate that the ^137^Cs activity concentrations in *A. elata* buds were lower at sites with less initial ^137^Cs deposited, regardless of the presence/absence of decontamination practices.

**Table 1.** Results of the linear mixed model used to test for effects of the dose rate at the soil surface, organic soil layer thickness, and slope steepness on the ^137^Cs activity concentrations in *Aralia elata* buds. Bold characters indicate statistical significance.

| Explanatory variable | Estimate | Standard error | *t* | *P* |
| --- | --- | --- | --- | --- |
| log<sub>e</sub> dose rate | 1.815 | 0.277 | 6.563 | **<0.001** |
| Organic soil layer thickness | 0.112 | 0.012 | 9.511 | 0.070 |
| Slope steepness | -0.021 | 0.017 | -1.222 | 0.223 |

**Fig 3.**
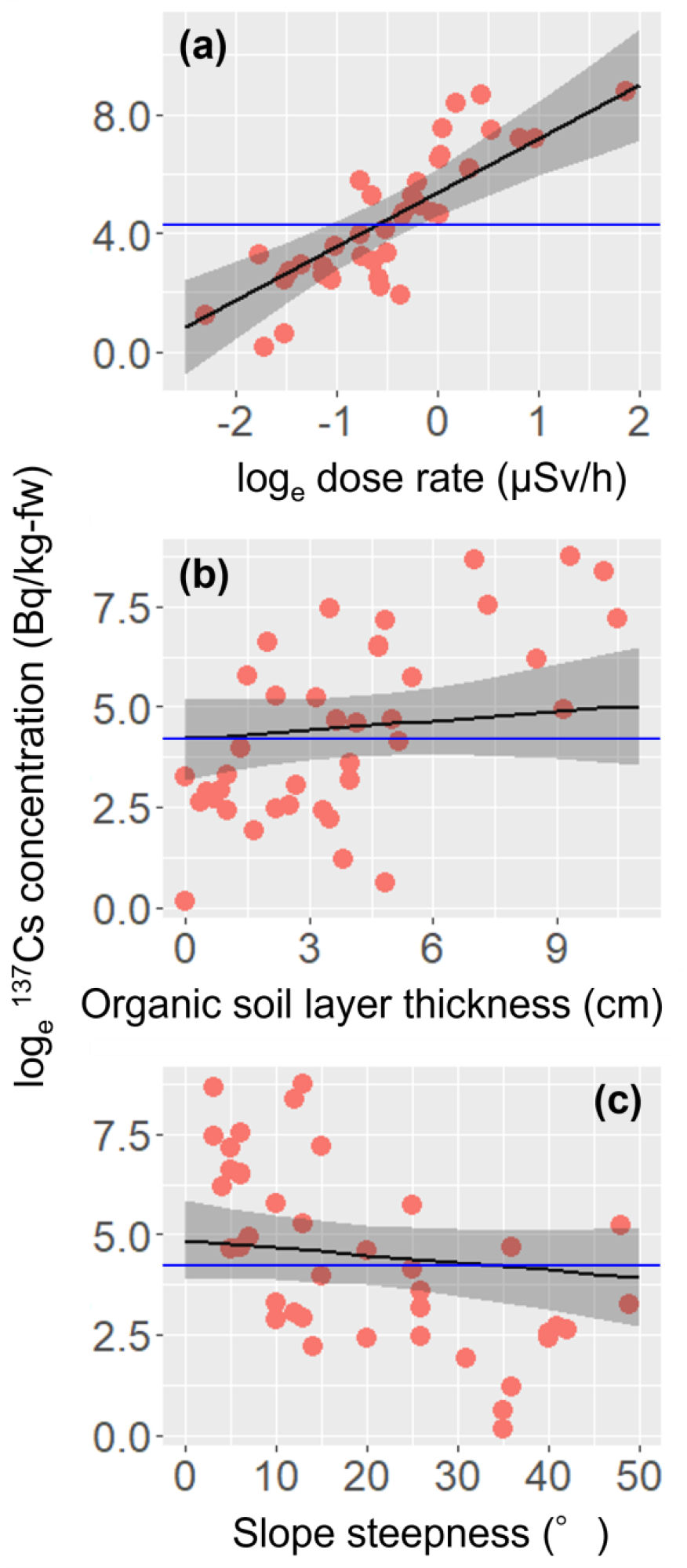
Relationships between ^137^Cs activity concentrations in *A. elata* buds and (a) the dose rate at the soil surface, (b) organic soil layer thickness, and (c) slope steepness. The blue horizontal lines indicate the Japanese reference level for food (100 Bq/kg). Regression lines were generated based on the results of the full linear mixed model. Explanatory variables other than those shown on the x axis were fixed at their mean values. Shaded bands indicate 95% confidence intervals.

**Fig 4.**
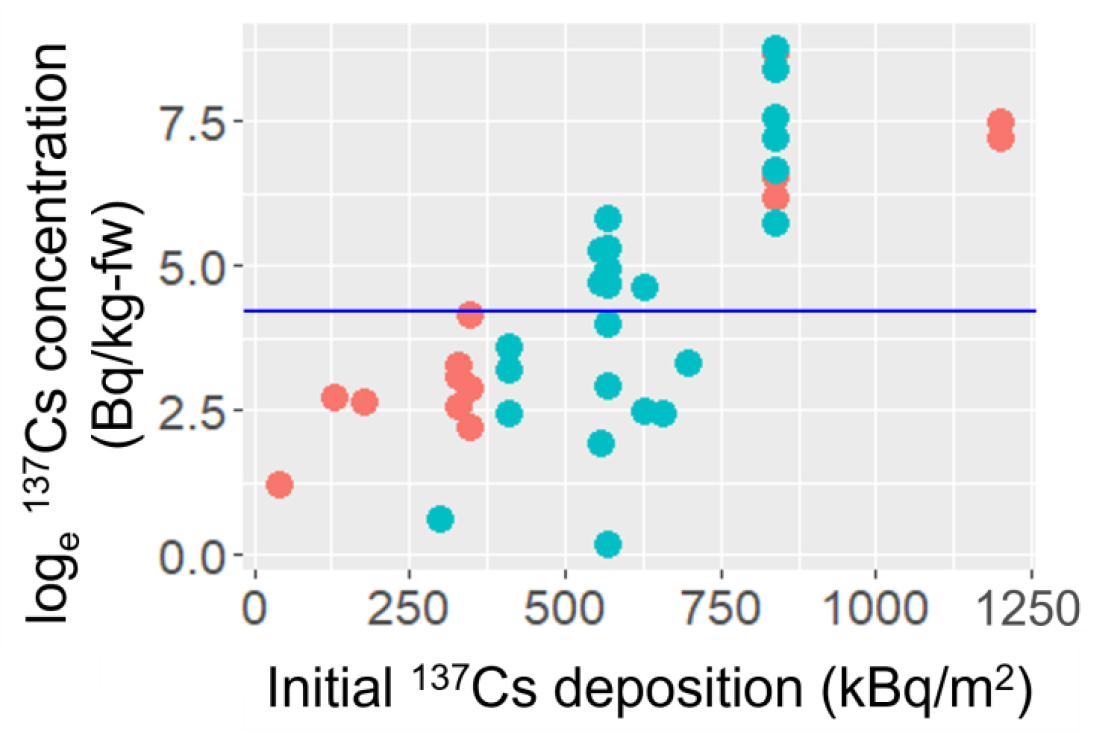
Relationship between ^137^Cs activity concentrations in *A. elata* buds and initial ^137^Cs depositions at the decontaminated (light blue) and abandoned (pink) sites. The blue horizontal line indicated the Japanese reference level for food (100 Bq/kg).

### Aggregated transfer factors

As previously reported [22], the ^137^Cs inventory in the soil and the dose rate at the soil surface were positively correlated (Fig 5). The relationship was expressed as Eq. 1: log_e_ (^137^Cs inventory in soil) = 1.23 × log_e_(dose rate on soil surface) + 6.03 (1) The slope and intercept were significant (*P* < 0.001) and the adjusted *R*^2^ was 0.856. The^137^Cs inventory in the soil estimated based on the relationship ranged from 24 to 4,200 kBq/m^2^ at the base of the *A. elata* plants. The estimated ^137^Cs inventory in the soil and ^137^Cs activity concentrations in *A. elata* buds were used to calculate the T_ag_ values, and the geometric mean (GM) and geometric standard deviation (GSD) of the T_ag_ were 3.7 × 10^−4^ m^2^/kg and 1.3, respectively (with a minimum of 2.3 × 10^−5^ m^2^/kg and a maximum of 8.5 × 10^−3^ m^2^/kg).

**Fig 5.**
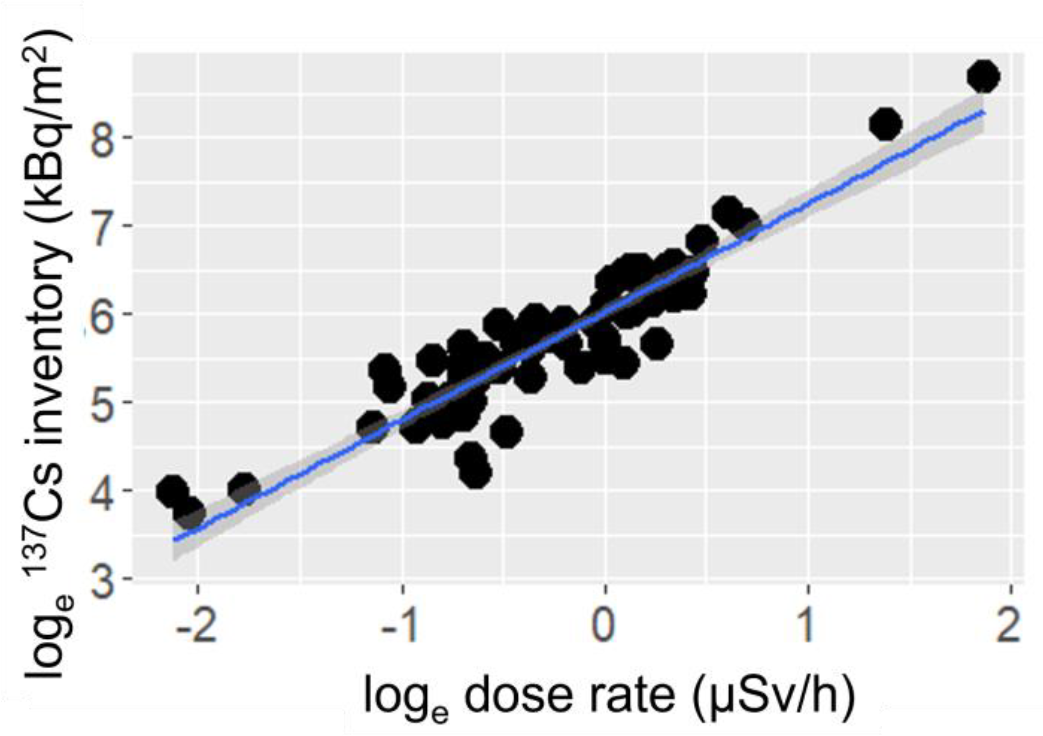
Relationship between the ^137^Cs inventory in the soil and the dose rate on the soil surface. Shaded bands indicate 95% confidence intervals.

## Discussion

The present study confirmed that ^137^Cs activity concentrations varied widely among *A. elata* buds, including concentrations above and below the Japanese reference level for food. In particular, plants at sites with a higher dose rate at the soil surface exhibited elevated ^137^Cs activity concentrations in their buds. These findings emphasize the importance of refraining from collecting and consuming *A. elata* buds from such habitats as a way to mitigate internal exposure. Interestingly, contrary to our expectations, decontamination practices did not lead to reduced ^137^Cs activity concentrations in the *A. elata* buds. This result further emphasizes the value of directly assessing the current dose rate at the soil surface at the base of *A. elata* plants rather than a decontamination record. This approach offers a practical means to avoid collecting buds with higher ^137^Cs concentrations.

Given that the dose rate at the soil surface was a useful indicator for estimating the ^137^Cs inventory in the soil (Eq. 1) [22], using a portable dosimeter was a convenient way to instantly estimate the ^137^Cs inventory in soil and associated transfer factors between the soil and organisms. The GM and GSD of the T_ag_ for *A. elata* buds were 3.7 × 10^−4^ m^2^/kg and 1.3 for 2022–2023, respectively. The GM and GSD values were smaller than those reported previously for *A. elata* buds (1.1 × 10^−3^ m^2^/kg and 3.1 for 2016–2018 [22] and 4.3 × 10^−4^ m^2^/kg and 4.0 for 2014–2019 [21]). The smaller GM value suggests a gradual reduction in the transfer of ^137^Cs from the soil to *A. elata* buds [27]. This decrease may be attributable to downward ^137^Cs migration in soils [28], followed by fixation in clay minerals, which subsequently decreases the bioavailability of ^137^Cs [29]. In addition, the remarkably smaller GSD value suggests that such a migration may induce a steady-like state of ^137^Cs dynamics in the soil, showing the clear positive relationship between the dose rate at the soil surface and ^137^Cs activity concentrations in *A. elata* buds. Although T_ag_ is generally rather variable (and thus often represented with GM and GSD) [8, 21, 22], the positive relationship may become observable under the current soil conditions more than 10 years after the contamination event.

The LMM demonstrated that the dose rate at the soil surface had a significant effect on ^137^Cs activity concentrations in *A. elata* buds. This finding agrees with previous studies that have reported positive relationships between the dose rate (or ^137^Cs inventories in the soil) and the ^137^Cs concentrations in various taxa [12–14]. The ^137^Cs inventory in the soil described herein was presumably affected by the amount of initial ^137^Cs deposited and by slope steepness, which is an important driver of soil runoff and affects the amount of ^137^Cs in soil [17]. However, the effect of steepness on concentrations in *A. elata* buds was minimal, suggesting that runoff of ^137^Cs may have been limited in the slopes of the *A. elata* habitats, where open sunlit environments often give rise to luxuriant vegetation that prevents soil erosion [30].

In addition, the thickness of the organic soil layer did not increase the ^137^Cs activity concentrations in *A. elata* buds, while the soil layer was assumed to possess abundant bioavailable ^137^Cs that can be transferred to plants [29]. For example, *Eleutherococcus sciadophylloides* (Kashiabura, the Queen of *sansai*), which develops extensive roots in the interface between the organic and mineral soil layers [31] exhibit one of the highest bud ^137^Cs concentrations among Japanese wild vegetables [6–8]. In addition, ^137^Cs activity concentrations in *E. sciadophylloides* are well correlated with ^137^Cs inventories in the organic soil layer [8]. Thus, ^137^Cs in the organic soil layer is still notable to understand ^137^Cs transfer from the soil to wild vegetables. However, the thickness of the layer may not always be correlated with the ^137^Cs inventory in it; thus, our result may not show a clear positive relationship between organic soil layer thickness and ^137^Cs activity concentrations in *A. elata* buds. Although the inventory in this layer is expected to be a good predictor of the ^137^Cs concentrations in *A. elata* buds, the estimate is generally time-consuming and is not a feasible way for the public and thus not fitted to the objective of this study.

The presence/absence of decontamination practices did not affect the ^137^Cs activity concentrations in *A. elata* buds. Although the specific processes leading to this result are unclear, it can be speculated from several processes. First, the decontamination practices were designed to decrease external exposure to less than 1 mSv/y (0.23 μSv/h) for local residents, and achieving this purpose in highly contaminated areas is a challenge [15]. Therefore, our sites may possess sufficient ^137^Cs in soils to have high ^137^Cs concentrations in *A. elata* buds even if decontamination practices were implemented. Second, the initial ^137^Cs deposition data we used [18] might be too coarse (250 m mesh) to detect the effect of decontamination on ^137^Cs activity concentrations in *A. elata* buds. Third, recontamination from surrounding environments might mask the potential effect of decontamination. For example, several studies have reported that soil surface removal as a decontamination practice does not effectively reduce the ^137^Cs concentrations in tree species [32, 33]. Moreover, recontamination from the surrounding environment can increase the levels in soils after decontamination [19]. The assessment of such effects is of utmost importance to mitigate external exposure and revive traditional practices of wild vegetable collection and consumption.

The present study revealed that ^137^Cs activity concentrations in *A. elata* buds increased as the dose rate at the soil surface increased. This environmental factor is easily measurable and helpful to avoid collecting *A. elata* buds with high ^137^Cs concentrations. Our findings indicate that the presence/absence of a decontamination practice did not affect the concentrations in *A. elata* buds. Further studies that examine the causality among the relationships between these environmental variables and ^137^Cs concentrations in *A. elata* are needed to understand the ^137^Cs dynamics between the soil and plants. Our findings provide an easy way to decrease the risk of collecting and consuming highly contaminated *A. elata* buds. Because cooking potentially decreases ^137^Cs concentrations in wild vegetables [34], a combination of collecting and cooking measures could further help revive this tradition. Developing measures to decrease the risk of collecting and consuming wild vegetables as reported here is now growing in significance to facilitate the revival of traditional cultures in polluted regions.

## Acknowledgements

We thank Mr. Kazuo Sasaki and Dr. Yoichi Tao for enabling us to collect samples, and the Ministry of the Environment for providing us with the data of presence/absence of decontamination practice at the study sites. This study was performed by the commissioned research fund by F-REI (JPFR23050301).

## Supporting information

**Table S1. The raw data of** ^**137**^**Cs activity concentrations in *Aralia elata* buds and environmental variables analyzed in Sakai et al. Simple ways to avoid collecting highly** ^**137**^**Cs-contaminaed *Aralia elata* buds**.

(XLSX)

## Author contributions

**Conceptualization:** Masaru Sakai, Mirai Watanabe, Seiji Hayashi

**Data curation:** Masaru Sakai, Mirai Watanabe, Masami K. Koshikawa, Asuka Tanaka, Akiko Takahashi, Seiichi Takechi, Masanori Tamaoki

**Formal analysis:** Masaru Sakai

**Funding acquisition:** Seiji Hayashi

**Investigation:** All authors

**Methodology:** Masaru Sakai

**Visualization:** Masaru Sakai

**Writing – original draft:** Masaru Sakai

**Writing – review & editing:** All authors

